# To sort or to hash? Revisiting *k*-mer counting

**DOI:** 10.64898/2026.11.23.768625

**Authors:** Victor Levallois, Rob Patro, Pierre Peterlongo, Giulio Ermanno Pibiri

## Abstract

**Motivation:** Counting *k*-mers in large genomic datasets is a foundational primitive in bioinformatics, supporting applications such as genome assembly, variant discovery, metagenomic profiling, and sequence database indexing. As sequencing throughput grows, *k*-mer counting is routinely applied to collections of hundreds to thousands of files. Even a modest per-file speedup compounds into hours of saved computation across such collections.

**Methods:** Sorting and hashing are the two main ways of counting *k*-mers, and practice has largely settled on the former, of which KMC3 [Kokot et al., 2017c] is the most refined expression. We reconsider this choice and analyze both approaches using a cost model that charges computation and memory transfers together. We show that the outcome of the comparison rests on two matters: how abundant the *k*-mers are, and whether the hash tables fit in cache. Sorting is faster only when data is mostly unique and the hash tables do not fit in cache.

We then design a *k*-mer counting algorithm in the light of this analysis. We adopt the consolidated paradigm based on first partitioning the *k*-mers by their minimizer, thus obtaining partitions of super-*k*-mers, and then counting each partition independently in parallel. In particular, super-*k*-mers are deduplicated before counting, and the number of partitions is chosen so that the hash tables fit in cache whenever possible. The algorithm avoids writing temporary data to disk if enough internal memory is available.

**Results:** We implemented this strategy in a tool named Tuna and show that it is faster than KMC3 on all but one of our inputs, spanning bacterial and human assemblies, metagenomes and read sets, by up to 4.3×. The advantage of Tuna compared to KMC3 in the counting step alone widens to 6.8× as *k*-mer abundance grows, as the analysis predicts. On the largest input tested, 614 GB of human sequencing reads, Tuna counts within 4 GB of memory and 0.7 TB of temporary disk. On that input the two tools finish within 1.1× of each other under the full 256 GB budget, and restricting every tool to the same 8 GB budget widens the advantage of Tuna over KMC3 to 1.8×.

**Software:** The software is freely available at github.com/vicLeva/tunaunder the MIT license.

## 1. Introduction

Counting the substrings of fixed length *k* (the *k*-mers) of a collection of DNA sequences is one of the most common preprocessing steps in bioinformatics. The result of this step associates each distinct *k*-mer with an integer, its frequency *count*, equal to its number of occurrences in the collection. These *k*-mer counts are central to genome assembly [Bankevich et al., 2012; Simpson et al., 2009] and assembly quality assessment [Rhie et al., 2020], and are stored explicitly by weighted *k*-mer dictionaries [Pibiri, 2023; Levallois et al., 2024]. They are also used when building indexes for pseudoalignment and sequence search [Alanko et al., 2023; Levallois et al., 2026; Lemane et al., 2024; Fan et al., 2024], to discard low-count *k*-mers that likely arise from sequencing errors.

As sequencing throughput grows, applications process collections of hundreds to thousands of genomes, with total input sizes reaching tens to hundreds of gigabytes. A fast and memory-efficient *k*-mer counting algorithm is therefore important to avoid limiting the throughput of the whole downstream pipeline.

Different *k*-mer counting algorithms have been proposed: some employ in-memory hash-table [Marçais and Kingsford, 2011; Melsted and Pritchard, 2011; Wang et al., 2021; Pandey et al., 2018]; others resort to disk-partitioned pipelines that exploit super-*k*-mers to reduce per-record I/O [Rizk et al., 2013; Li et al., 2015; Deorowicz et al., 2015; Kokot et al., 2017c], with recent work exploring more compact representations that exploit the *k* − 1 base overlap between consecutive super-*k*-mers, suited to large *k* and long-read data [Martayan et al., 2025], or targeting massively parallel hardware [de Oliveira and Fernandes, 2026]. Underlying these tools sits a structural choice between sorting and hashing a partition of *k*-mers, and the state-of-the-art tool, KMC3 [Deorowicz et al., 2015; Kokot et al., 2017c], is based on sorting. This paper revisits the comparison between sorting and hashing as fundamental paradigms to solve the *k*-mer counting problem.

### Our contribution

We give a formal analysis of the two strategies, sorting and hashing, under different computational models: the classic RAM model, the two-level I/O model [Vitter, 2001], and a cost model that charges CPU work and memory transfers together (Section 4). The analysis identifies two quantities that determine which strategy prevails: the average abundance of the *k*-mers, which we use as a proxy for the coverage of the input, and whether the hash tables employed by the hashing strategy remain resident in cache. Sorting is faster than hashing only when: 1. the data is largely free of repetitions, and 2. the hash tables used by the hashing strategy exceed the cache.

We develop a *k*-mer counting algorithm informed by this analysis (Section 5). We follow the established paradigm of partitioning the *k*-mers according to their minimizers [Schleimer et al., 2003; Roberts et al., 2004], which yields partitions of super-*k*-mers that are subsequently counted independently and in parallel. Two design decisions follow from the analysis: super-*k*-mers are deduplicated before counting, and the partitions are sized so that the hash tables remain within the cache. Provided that sufficient internal memory is available, the algorithm writes no temporary data to disk.

We implemented Tuna, a C++ implementation of this hash-based approach, built on engineered primitives that have recently matured: SIMD-accelerated sequence parsing [Martayan et al., 2026] and a high-performance hash table [Leitner, 2022] (Section 5).

We compare Tuna against FastK [Myers, 2021] and KMC3 [Kokot et al., 2017c], the latter being the *de-facto* algorithm for *k*-mer counting. We show that Tuna is faster than KMC3 on all but one of our inputs, spanning bacterial and human assemblies, metagenomes and read sets, by up to 4.3×, and faster than FastK by up to 3.8×. In the counting phase alone, its advantage widens to 6.8× as the abundance of the *k*-mers grows, as the analysis predicts. On the largest input we consider, 614 GB of human sequencing reads, Tuna runs within 4 GB of memory and 0.7 TB of temporary disk, its disk pipeline taking over once the partitioned data outgrows the memory budget. Under the full 256 GB budget, it is 1.1× faster than KMC3 there, and restricting every tool to the same 8 GB budget widens that to 1.8×.

## 2. Background and problem definition

This section summarizes basic background information and formalizes the *k*-mer counting problem.

Since DNA is double-stranded, a *k*-mer *x* and its reverse complement rc(*x*) represent the same genomic string on opposite strands. The *canonical* form of a *k*-mer *x* is canon(*x*) = min(*x*, rc(*x*)) under lexicographic order. In the following, let *S* be a hypothetical string over the DNA alphabet ∑.

### Definition 1

(Minimizer). *Given m, k* ∈ N, *m < k, and a total order* O *over all m-mers, the minimizer of a k-mer is its smallest m-mer under* O. *The canonical minimizer of a k-mer x, µ*(*x*), *is the smallest m-mer in* {canon(*y*) : *y m*-mer of *x*} *under* O.

Some remarks are in order. Note that according to the above definition, a minimizer is an *m*-mer, not an occurrence: repeated occurrences of the smallest *m*-mer in the *k*-mer leave the minimizer unchanged, so no tie-breaking rule is needed. Canonical minimizers are *strand-invariant*, i.e., a *k*-mer *x* and rc(*x*) have the same canonical minimizer because the *m*-mers of rc(*x*) are the reverse complements of the *m*-mers of *x*. In practice, O is instantiated as a pseudo-random hash function, yielding the *random* minimizer scheme. We use canonical random minimizers for the rest of the paper.

One important property of minimizers is that consecutive *k*-mers of *S* tend to share the same minimizer, as the minimizer changes only when the sliding window “loses” its current minimum from the left or “gains” a new one from the right. Under a random order, the expected fraction of *k*-mer positions at which the minimizer changes is ≈ 2*/*(*k* − *m* + 2) [Schleimer et al., 2003] (this holds for the canonical minimizer too but its proof is omitted for brevity).

### Definition 2

(Super-*k*-mer). *A* super-*k*-mer *is a maximal run of consecutive k-mers of S that share the same minimizer*.

The length of a super-*k*-mer is between *k* and 2*k* − *m* bases. A super-*k*-mer of length *l* encodes *l* − *k* + 1 consecutive *k*-mers in *l* bases, whereas storing *k*-mers individually would repeat *k* − 1 bases between each consecutive pair.

### Problem 1

(Canonical *k*-mer counting). *Given a collection S of DNA strings and an integer k >* 0, *let f* : ∑^*k*^ → ℕ_0_ *where f* (*x*) = ∑_*S∈S*_ |{*j* : canon(*S*[*j* .. *j* + *k*)) = *x*}| *and f* (*x*) = 0 *for any x not present in S. The* canonical *k*-mer counting *problem is to produce the set* {(*x, f* (*x*)) : *c*_min_ ≤ *f* (*x*) ≤ *c*_max_} *for user-supplied thresholds c*_min_, *c*_max_ ∈ ℕ.

Setting *c*_min_ *>* 1 suppresses *k*-mers likely arising from sequencing errors, while *c*_max_ caps highly repetitive *k*-mers.

## 3. Related work

Two main families of *k*-mer counting algorithms have been proposed in the literature, based on either hashing or sorting. Most of these algorithms share the same disk-based organization, so we describe it once for both families. Every super-*k*-mer of *S* is assigned to a partition determined by hashing its minimizer. Since a *k*-mer and its reverse complement have the same canonical minimizer, all occurrences of a *k*-mer, on either strand, fall in the same partition. Each partition can therefore be counted independently, with no synchronization between partitions and no final merging step. We refer throughout to the two resulting phases as Phase 1, which streams the input and builds the partitions, and Phase 2, which computes the counts of a partition.

### Hashing

The first challenge addressed by hash-based counters was parallelism, that is, allowing many threads to insert into a single shared table without locks. Jellyfish [Marçais and Kingsford, 2011] achieves this by having each thread acquire a slot with a single atomic instruction, retrying upon failure, which yields near-linear scaling with the number of cores. Its table, however, must be sized before execution begins; if the estimate is too low, the overflow is written to disk and merged at the end, a recovery mechanism rather than a design intended for data larger than memory. Reducing peak memory was the next concern. BFCounter [Melsted and Pritchard, 2011] filters erroneous singletons through a Bloom filter and inserts only confirmed *k*-mers into the table, while Squeakr [Pandey et al., 2018] replaces the table itself with the Counting Quotient Filter [Pandey et al., 2017], a compact structure that supports exact and approximate modes and an on-disk representation. Cache efficiency under frequent collisions motivated the lock-free chaining table of CHTKC [Wang et al., 2021]. Handling datasets larger than the available memory required a different organization. DSK [Rizk et al., 2013] distributes the *k*-mers into *n* partitions on disk and counts each partition in a separate hash table, so that peak memory is *O*(input size*/n*) regardless of the dataset. The disk-based pipeline it introduced has since been adopted by most counters, including the one presented in this paper. Subsequent work reduced the size of the partitions themselves: MSPKmerCounter [Li et al., 2015] was the first counter to employ minimizers, encoding consecutive *k*-mers that share a minimizer as a single record (a super-*k*-mer) rather than as separate records. The super-*k*-mer representation was subsequently adopted by both families.

### Sorting

The sorting-based approach was popularized by KMC2 [Deorowicz et al., 2015], which combined super-*k*-mer partitioning with a least-significant-digit radix sort and became the *de facto* standard disk-based counter. Since this sorting algorithm requires many passes on long keys, KMC3 [Kokot et al., 2017c] replaced it with a most-significant-digit radix sort, together with further I/O improvements and a set of operations over *k*-mer databases. The same approach is followed by FastK [Myers, 2021], which has not appeared as a peer-reviewed publication but is often used as the counter in the genome-profiling step of recent long-read assembly pipelines [Wright et al., 2024; Bein et al., 2025; Xing et al., 2025]. Sorting is also used by Meryl, the counting component of the Merqury assembly-quality framework [Rhie et al., 2020], which provides a rich algebra of *k*-mer set operations intended for comparing assemblies rather than for fast counting.

### Specialised counters

Other designs target more specific settings or hardware. Assembled genomes, whose *k*-mer diversity is low, are the focus of KCOSS [Tang et al., 2022], which combines a segmented Bloom filter with cuckoo hashing; its design is, however, so closely tailored to this input that it is poorly suited to the general multi-file setting. KCMBT [Mamun et al., 2016] explores burst trees as a cache-aware alternative to both hashing and sorting. The growth of memory with *k* is addressed by KFC [Martayan et al., 2025], whose *hyper-k-mers* merge consecutive super-*k*-mers through their overlap of *k* − 1 bases, lowering the asymptotic bound from 6 to 4 bits per nucleotide; it is the first counter whose memory scales sub-linearly with *k*, a clear advantage for large *k*. Gerbil [Erbert et al., 2017] and RapidGKC [Cheng et al., 2024] follow the usual two-phase organization but achieve their best throughput only on machines equipped with a GPU. Finally, BIKE [de Oliveira and Fernandes, 2026] formulates counting as a binary matrix of pairwise character comparisons over the input, two *k*-mers being identical when all *k* comparisons agree. Its memory footprint depends only on the sequence length, and not on the alphabet or on *k*, which is valuable for amino acid sequences with large *k*; it targets massively parallel hardware such as FPGAs, and its authors present the current implementation as a proof of concept rather than a production tool.

## 4. To sort or to hash?

The starting point for our development is KMC3 [Kokot et al., 2017c], the current state of the art for the *k*-mer counting problem. Like several other approaches reviewed in Section 3, KMC3 first partitions the input into sets of super-*k*-mers, and subsequently determines, for each set, the distinct *k*-mers it holds together with their abundances using sorting. The present section analyses this second step (where two different paradigms can be adopted) to address a basic question: for a given partition of super-*k*-mers, how does hashing perform instead of sorting?

The question is addressed through the analysis of two abstract algorithms. Algorithm S is intended to model the specific sort-based approach adopted by KMC3 as faithfully as possible (not *any* sort-based approach). Algorithm H, instead, first deduplicates the set of super-*k*-mers using a hash table and then inserts the *k*-mers contained in the deduplicated set into another hash table to count them.

### Computational models

Throughout the analysis, we use two computational models, the RAM model and the two-level I/O model [Vitter, 2001]. In the RAM model, every machine operation and every memory access costs one unit of work, and the cost of an algorithm is the number of operations it executes. In the two-level I/O model, the machine has a fast memory (cache) of *M* bytes and a slow memory of infinite size, and data moves between the two in blocks (cache lines) of *B* bytes. The cost of an algorithm in this model is the number of block transfers, and computation on data residing in fast memory is free.

The two models are deliberately complementary: the RAM model charges work and ignores different memory devices, the I/O model charges data movement and ignores work. We will therefore combine them into a single cost model, charging both CPU work and memory transfers, to obtain a more realistic model.

### Notation and setting

All quantities refer to a single partition. The partition holds *u* super-*k*-mers containing *N k*-mer occurrences in total, so that *L* = *N/u* is the average super-*k*-mer length, measured in *k*-mers. The number of distinct *k*-mers in the partition is *d*, and *c* = *N/d* is the average abundance, which we use as a proxy for coverage. Among the *u* super-*k*-mers, *u*_*d*_ are distinct, and *N*_*d*_ denotes the number of *k*-mer occurrences contained in the distinct super-*k*-mers, so that *d* ≤ *N*_*d*_ ≤ *N*. We assume *k*-mers over an alphabet of size four, so each base is coded in two bits. A super-*k*-mer of *j k*-mers occupies *j* + *k* − 1 bases, so the packed partition holds *N* +*u*(*k*−1) = *N* (1+(*k*−1)*/L*) bases, that is *N* (1+(*k*−1)*/L*)*/*4 bytes. Each entry of the output (*k*-mer plus counter) occupies *s*_out_ bytes. Both algorithms read the packed partition and write the output. We indicate with *R* the number of sequential block transfers issued during the read of the input and with *W* the ones issued during the write of the output^1^:

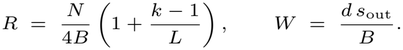

Both are charged identically to the two algorithms.

### 4.1. The sort-based algorithm S

Algorithm S consists of four stages: expand, sort, compact, and merge.

**Expand**. S does not sort *k*-mers but (*k, x*)-mers [Deorowicz et al., 2015], with *x* = min(31−(*k* mod 32), 3) if (*k* mod 32)≠ 0 and *x* = 0 otherwise: records of up to *k* + *x* symbols, each covering up to *x* + 1 consecutive *k*-mers, obtained by tiling every super-*k*-mer with cuts *x* + 1 positions apart. The number of *k*-mers a record covers beyond the first, called its *extension x*′ ∈ {0, …, *x*}, is *x* for all records except possibly the last one of each super-*k*-mer. A super-*k*-mer of *l* bases holds *l* − *k* + 1

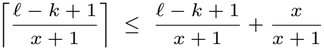

*k*-mers and thus yields

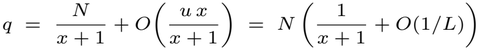

records, and summing over all super-*k*-mers in the partition, the partition has records in total, of which *q*_*d*_ are distinct. A record is stored in whole 64-bit words and occupies *s* = 8⌈(2(*k* + *x*) + 2)*/*64⌉ bytes, the two extra bits storing the extension *x*′. Since a record covers at most *x* + 1 distinct *k*-mers, *q*_*d*_ ≥ *d/*(*x* + 1). Since the records produced from duplicate super-*k*-mers coincide and the distinct records are among those produced from the distinct super-*k*-mers, we have *q*_*d*_ ≤ *N*_*d*_(1*/*(*x* + 1) + *O*(1*/L*)).

The expansion itself is one pass over the packed partition producing the *q* records: in the RAM model this costs Θ(*N*), as every base is decoded once, while in the I/O model the stage reads the packed partition and writes the records, *R* + *qs/B* blocks, all sequentially.

### Sort

The sorting stage uses a most-significant-digit (MSD) radix sort algorithm over the *q* records of (*k, x*)-mers. This algorithm models the RADULS [Kokot et al., 2017a,b] algorithm used by KMC3. Each record has *P* = ⌈(2(*k*+*x*)+2)*/r*⌉ digits of *r* bits each. The algorithm has the following features.

- A small-sort threshold *θ*: a bucket of at most *θ* records is sorted with a comparison sort instead of being split further. For example, Kokot et al. [2017a] sets *θ* = 384 experimentally.
- No test for homogeneous buckets: a bucket of all-equal keys is split recursively anyway.
- A level whose buckets exceed the cache performs one sequential read and one sequential write of its data.

The work of the stage depends on how the keys distribute among the buckets, and we analyse two cases, both of which occur in practice: the worst case, and the case of keys distributing near-uniformly across the 2^*r*^ buckets of a level.

In the worst case, a record is inspected at most once per key digit, that is at most *P* times, and takes part in at most one comparison sort, which adds at most *O*(log *θ*) comparisons per record: *O*(*q* (*P* + log *θ*)) work in the worst case. This is the regime of highly repetitive inputs, where a bucket of one key repeated more than *θ* times is split at every level without ever shrinking below the threshold.

For near-uniform keys, instead, the recursion has height 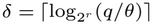, buckets leave it with Θ(*θ*) records, and the work is about Θ(*q* (*δ* + log *θ*)). This is the regime of partitions of mostly distinct records: the records of a partition share their minimizer, but the remaining bases vary, so the leading digits spread across the buckets.

Either way the cost depends on *q* only, never on *q*_*d*_: the sort is oblivious to duplicates.

In the I/O model, a level whose buckets exceed the cache reads every record once and writes it once to its destination bucket, i.e. one full sequential read and one full sequential write of the array, 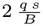 transfers, and no random accesses occur. Near-uniform keys have 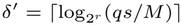 such levels, and in the worst case there are up to *P*, again under heavy duplication, a homogeneous bucket larger than the cache surviving every level. In total, the sorting stage costs Θ(*q* (*δ* + log *θ*)) work and 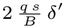 sequential transfers for near-uniform keys, and *O*(*q* (*P* + log *θ*)) work and 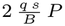 transfers in the worst case.

### Compact

A linear scan collapses runs of identical records into one record with a counter, leaving *q*_*d*_ records: Θ(*q*) in the RAM model, at most 2*qs/B* sequential transfers in the I/O model (the array is read once and the compacted records are written once).

### Merge

The sorting stage operates on (*k, x*)-mer records, not on *k*-mers, and the same *k*-mer sits in up to *x* + 1 records at different offsets, far apart in sorted order, so a final merge reassembles the counts [Deorowicz et al., 2015]. Records with equal extension *x*′ and equal first *z* symbols (0 ≤ *z* ≤ *x*′) form a contiguous range of the sorted array, and the *k*-mers at offset *z* within that range appear in sorted order. There is one such implicit sorted list per (*x*′, *z*, prefix), i.e. 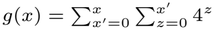 lists, for example *g*(*x*) = 6, 27, 112 for *x* = 1, 2, 3, respectively. A binary min-heap over the non-empty lists emits the sorted *k*-mer stream, summing counters of equal *k*-mers. The number of *k*-mer items it processes is *η* = (*x*′ + 1) over the distinct records, which satisfies *d* ≤ *η* ≤ *N*_*d*_: every distinct *k*-mer occurs in at least one distinct record and thus contributes at least one item, and the tilings of the distinct super-*k*-mers partition their *N*_*d*_ *k*-mer occurrences.

The merge thus costs Θ(*η*) in the RAM model (*η* heap operations of log_2_ *g*(*x*) = Θ(1) comparisons each), between Θ(*d*) and Θ(*N*_*d*_).

In the I/O model, it costs *q*_*d*_*s/B* + *W* sequential transfers, the heap and the list cursors being cache-resident. For *x* = 0 every record is a *k*-mer: no merge is needed, and the compaction already emits the output.

#### Proposition 4.1.

*The cost of algorithm* S *is*

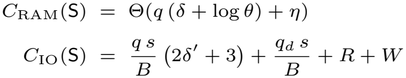

where *d*≥ *N*_*d*_≥*N*.

*for near-uniform keys, and with δ and δ*′ *both replaced by P in the worst case. The value of q is N* (1*/*(*x*+1)+*O*(1*/L*)) *and η Is the number of k-mers processed by the merge, N/c* ≤ *η* ≤ *N*_*d*_.

Note that both costs in Proposition 4.1 are functions of the number of occurrences *N* and are insensitive to the abundance *c* at fixed *N*, except for the merge term Θ(*η*), which lies between Θ(*N/c*) and Θ(*N*_*d*_). All transfers are sequential.

### 4.2. The hash-based algorithm H

Algorithm H consists of three stages instead: deduplicate, count, output. The first two stages rely on hashing. For the analysis, we assume a hash table with *O*(1) expected insert and lookup time under the usual *simple uniform hashing* assumption. For a hash table of footprint *F* bytes probed at uniformly random positions, we write miss(*F*) = max(0, 1 − *M/F*) for the probability that a probe misses the cache (only an *M/F* fraction of the table is resident in cache).

#### Deduplicate

The *u* super-*k*-mers are inserted into a hash table keyed by super-*k*-mer, yielding the *u*_*d*_ distinct super-*k*-mers with multiplicities. Since super-*k*-mers have variable length, we charge the table by its total footprint *F*, the memory in bytes it occupies for the *u*_*d*_ distinct super-*k*-mers. The stage costs expected Θ(*u*) = Θ(*N/L*) in the RAM model (a super-*k*-mer, like a *k*-mer, occupies Θ(1) machine words for the *k* and *L* of interest), and in the I/O model *u* · miss(*F*) random block transfers plus the *R* sequential ones for reading the partition.

#### Count

Each distinct super-*k*-mer is expanded and its *k*-mers are inserted into a hash table of Θ(*d*) entries, with *s*_h_ bytes per entry, each insertion adding the multiplicity of the super-*k*-mer to the *k*-mer’s counter: *N*_*d*_ insertions, expected Θ(*N*_*d*_) RAM cost and *N*_*d*_ · miss(*ds*_h_) random block transfers.

#### Output

A scan of the table emits the (distinct *k*-mer, count) pairs: Θ(*d*) RAM cost and *W* sequential transfers.

##### Proposition 4.2.

*The cost of algorithm* H *is*

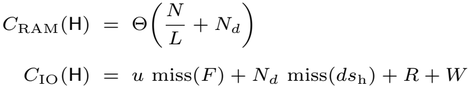

*where d* ≤ *N*_*d*_ ≤ *N*.

Note that both costs in Proposition 4.2 are governed by the numbers of *distinct* objects: the RAM cost through *N*_*d*_, the I/O cost through the table footprints *F* and *ds*_h_, so both decrease, per occurrence, as the abundance *c* grows and deduplication becomes effective. The miss(·) terms count *random* transfers.

### 4.3. Discussion and combined cost

We now compare the two algorithms using both computational models. In the RAM model, substituting *q* ≈ *N/*(*x* + 1) in Proposition 4.1 and dividing by the cost of Proposition 4.2, we obtain the ratio

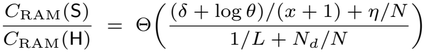

(with *P* in place of *δ* in the worst case). At *c* = 1 every record is distinct, *η* = *N*_*d*_ = *N*, and the ratio is 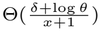. As *c* increases and deduplication becomes effective, *N*_*d*_*/N* → 0, and *η/N* → 0 with it since *η* ≤ *N*_*d*_, so the ratio grows by a factor of *L*, to 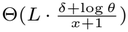 the RAM model predicts that H wins at every coverage, by a growing factor.

In the I/O model, the common transfers *R* and *W* are identical for the two algorithms, and we drop them from the comparison. What distinguishes the algorithms is the streaming of the records for S, (2*δ*′ + 3) *qs/B* + *q*_*d*_*s/B* sequential transfers (with *P* in place of *δ*′ in the worst case), against the table probes for H, *u* · miss(*F*) + *N*_*d*_ · miss(*ds*_h_) random ones.

At low coverage, *c* ≈ 1 and every record and every super-*k*-mer is almost distinct, i.e., *q*_*d*_ ≈ *q* and *N*_*d*_ ≈ *N*. Using again *q* ≈ *N/*(*x* + 1) and *u* = *N/L*, the comparison is between 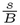· 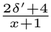 *sequential* I/Os per occurrence and:

- 1 + 1*/L random* I/Os when the tables do not fit in cache. In this case, H performs fewer I/Os only if

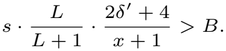

This fails, for example, for small *s* and *x >* 0 but can hold when *x* = 0 and records span multiple words, e.g., *k* = Under the parameter values of our experimental setup (Section 6), however, the inequality does not hold: the I/O model predicts that S wins over H in this scenario.
- no I/Os at all when the tables fit in cache. In this case, H outperforms S. Now let the coverage grow, at a fixed set of distinct *k*-mers. Both *q*_*d*_ and *N*_*d*_ count distinct super-*k*-mers, so they stop growing once every super-*k*-mer has been seen. The merge read of S, *q*_*d*_*s/B*, and the counting probes of H, *N*_*d*_ · miss(*ds*_h_), therefore stay bounded as *N* grows, and become negligible relative to the two costs that grow with *N* : S still passes over all *q* records, and H still probes its first table once per super-*k*-mer occurrence, *u* = *N/L* times. The comparison becomes (2*δ*′+3) *qs/B* sequential transfers against *N/L*·miss(*F*) random ones, and we have two regimes as before.
- The table exceeds the cache, the case on large workloads with large *d*: every probe misses, so H still pays *N/L* random transfers, a cost that no longer decreases with the coverage. Comparing it with the (2*δ*′ + 3) *qs/B* sequential transfers of S, and using *q* ≈ *N/*(*x* + 1), the algorithm H performs fewer I/Os only if

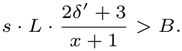

This holds for the usual parameters, and S stays ahead only for short super-*k*-mers (small *L*) and large *x*.
- The distinct set is small enough that the tables fit in the cache: the probing costs vanish for H, whereas S still pays its record passes, (2*δ*′ + 3) *qs/B*. In this case, H wins over S.

In summary, the I/O model predicts that H almost always wins except at low coverage when the hash tables do not fit in cache.

These comparisons, however, count I/Os of different kinds, since the model charges the same cost for a *sequential* transfer and for a *random* transfer. This is what the combined model in the next paragraph corrects.

#### Combined cost

We now model each elementary operation and each transfer with a constant, a proxy for the actual time on the target machine: *a*_*r*_ for one record-level step of the sort, either extracting a digit and writing the record to its bucket, or one comparison in the final small sorts, *a*_*c*_ for comparing one record with the previous one during the compaction, *a*_*m*_ for one comparison in the merge heap, *a*_*h*_ for a hash probe hitting the cache, *τ* for a sequential block transfer, and *λ* for a random cache miss. Summing the CPU and I/O costs of Propositions 4.1 and 4.2 weighted by these constants, and dividing by *N*, we obtain the cost per *k*-mer occurrence, in the near-uniform case (*P* replaces *δ* and *δ*′ in the worst case):

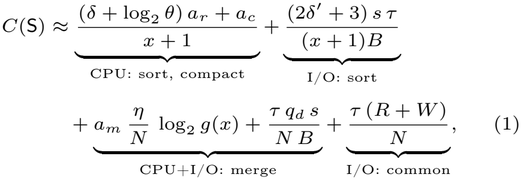

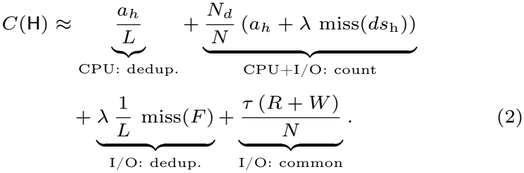

Equations (1) and (2) highlight what happens when the coverage grows, that is, *N* → ∞ while the set of distinct objects stays fixed: the two algorithms approach different constant floors. Writing *A* = (*δ* + log_2_ *θ*) *a*_*r*_ + *a*_*c*_ + (2*δ*′ + 3)*sτ/B* for the 6 To sort or to hash? Revisiting *k*-mer counting full per-record cost of S, its floor is *A/*(*x*+1): the idea of sorting (*k, x*)-mers spreads that cost over *x* + 1 occurrences, lowering the floor by a constant factor, at most 4 since *x* ∈ {0, 1, 2, 3}. The floor of H is instead (*a*_*h*_ + *λ* miss(*F*)) */L*, one probe per super-*k*-mer occurrence, charged a cache miss when the table stays out of cache. The floor of H is thus the lower of the two whenever *L A >* (*x* + 1) (*a*_*h*_ + *λ* miss(*F*)), which holds for the values of interest: S pays a full record-level cost every *x* + 1 occurrences, while H pays one probe every *L*.

#### Examples

We instantiate the two levels of the I/O model as the last-level cache and main memory: *M* is the size of the last-level cache and *B* = 64 bytes is a cache line, so *τ* is the amortized time of a sequential line transfer from memory, bandwidth-bound, and *λ* that of a random one, latency-bound. Representative values on one core of a modern machine are *a*_*r*_ ≈ *a*_*c*_ ≈ 1 ns, *a*_*m*_ ≈ 2 ns, *a*_*h*_ ≈ 5 − 10 ns, *τ* ≈ 3 ns. The value of *λ* deserves care: the latency of a memory access is 60 − 100 ns, but independent misses overlap (memory-level parallelism), and hash insertions of a key stream are independent, so their amortized cost per miss is 10 − 15 ns. Only dependent chains, as in pointer chasing, pay the full latency.

Consider the instantiation *r* = 8, *θ* = 384, *s* = 8, *s*_out_ = 12, *δ* = 2, *δ*′ = 1, as for partitions of a few hundred thousand records, *L* = 10, *a*_*h*_ = 7.5 ns and *λ* = 10 ns, for both *k* = 27 (*x* = 3) and *k* = 31 (*x* = 0: the merge disappears and the sort processes all *N* records). At *c* = 1 we have *η* = *N*_*d*_ = *N*; at *c* = 20 we take *N*_*d*_*/d* = 1.5, with *η* at its upper end *N*_*d*_ and *q*_*d*_ at its upper end *N*_*d*_*/*(*x* + 1).

The cost of S depends on *k* but not on the cache regime, since all its transfers are sequential. Equation (1) gives, in ns per occurrence: for *k* = 27, *C*(S) ≈ 18 at *c* = 1, decreasing to *C*(S) ≈ 4.5 at *c* = 20 as the merge term decays; for *k* = 31, *C*(S) ≈ 14 at *c* = 1 and *C*(S) ≈ 13.5 at *c* = 20, essentially flat, as there is no merge. The cost of H instead does not depend on *k* but on the cache regime. Equation (2) gives *C*(H) ≈ 20 at *c* = 1 and *C*(H) ≈ 3.1 at *c* = 20 when the tables do not fit in the cache (the random misses dominate), and *C*(H) ≈ 9 and *C*(H) ≈ 1.4, respectively, when they do, as for a partition with a small distinct set.

Comparing the two: when the tables do not fit in the cache, S is ahead at *c* = 1 (18 and 14 against 20), but the verdict reverses at *c* = 20 (3.1 against 4.5 and 13.5), with H ahead by a factor between 1.4 and about four. When the tables fit, H is ahead at every coverage, by a factor of three (*k* = 27) to an order of magnitude (*k* = 31) at *c* = 20.

#### Conclusions

Whenever the hash tables stay in the cache, algorithm H wins over S at every coverage, since a cached probe costs less than the record-level work of the sort. On the other hand, when the hash tables do not fit in cache, algorithm S wins on mostly unique data because sequential passes beat random misses. Algorithm H prevails as the data grows redundant, because super-*k*-mer deduplication shrinks its work with the abundance, while the sorted volume never shrinks. We remark again that this comparison is specific to the sorting strategy of KMC3, the state-of-the-art algorithm.

One may wonder whether a version of KMC3 modified in the light of these conclusions would have been faster. To this end, let S′ be the variant of S that deduplicates the super-*k*-mers exactly as H does and then sorts only the distinct records, each carrying a multiplicity (this raises *s* to *s*′). The cost of S′ is

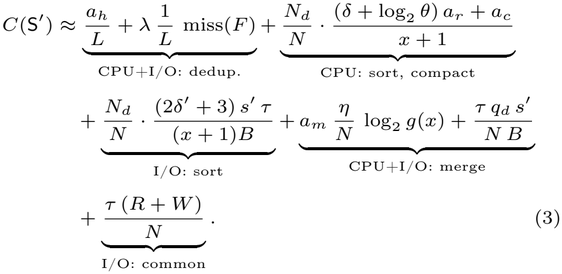

Comparing Equations (3) and (2), the “dedup” and “common” terms coincide, so the comparison is decided by the remaining terms. Note that each term carries a factor *N*_*d*_*/N* : the “sort” and “compact” terms explicitly, the “merge” terms after writing *η/N* = *N*_*d*_*/N* · *η/N*_*d*_ and *q*_*d*_*/N* = *N*_*d*_*/N* · *q*_*d*_*/N*_*d*_. Simplifying that common factor, the two algorithms are compared per *distinct* occurrence: S′ pays

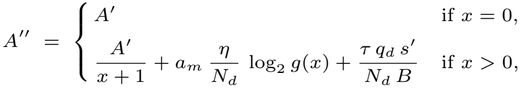

where *A*′ is *A* with *s*′ in place of *s*, while H pays one insertion into the *k*-mer hash table, i.e., *a*_*h*_ + *λ* miss(*ds*_h_). When the *k*-mer hash table fits in the cache, the insertion costs *a*_*h*_ alone, less than what S′ pays for the values of interest: H is faster. When it does not, H is faster whenever *A*″ *> a*_*h*_ + *λ* miss(*ds*_h_). The latter condition does not hold (and S′ is therefore faster) only when the *k*-mer hash table exceeds the cache widely enough that most probes miss.

Lastly, observe that when the coverage grows, i.e., *N* → ∞ while *N*_*d*_ stays fixed, every term carrying the factor *N*_*d*_*/N* vanishes and both algorithms approach the same constant floor: (*a*_*h*_ + *λ* miss(*F*))*/L*, the cost of deduplication.

## 5. Implementation

We implemented a *k*-mer counter in C++, named Tuna, whose counting phase (Phase 2) follows algorithm H (hashing-based) of Section 4, while Phase 1 is the partitioning step common to most counters (Section 3). Since the counting phase of KMC3 follows S (sorting-based), comparing the two tools tests the analysis empirically. This section describes the implementation of Tuna.

Figure 1 gives an overview of the algorithm. Phase 1 streams the input and routes every super-*k*-mer to a partition. Phase 2 then processes the partitions one at a time, counting the *k*-mers of each partition in a hash table. Running the two phases one after the other ensures that Phase 2 sees complete partitions, so counting needs no synchronization.

**Fig. 1.**
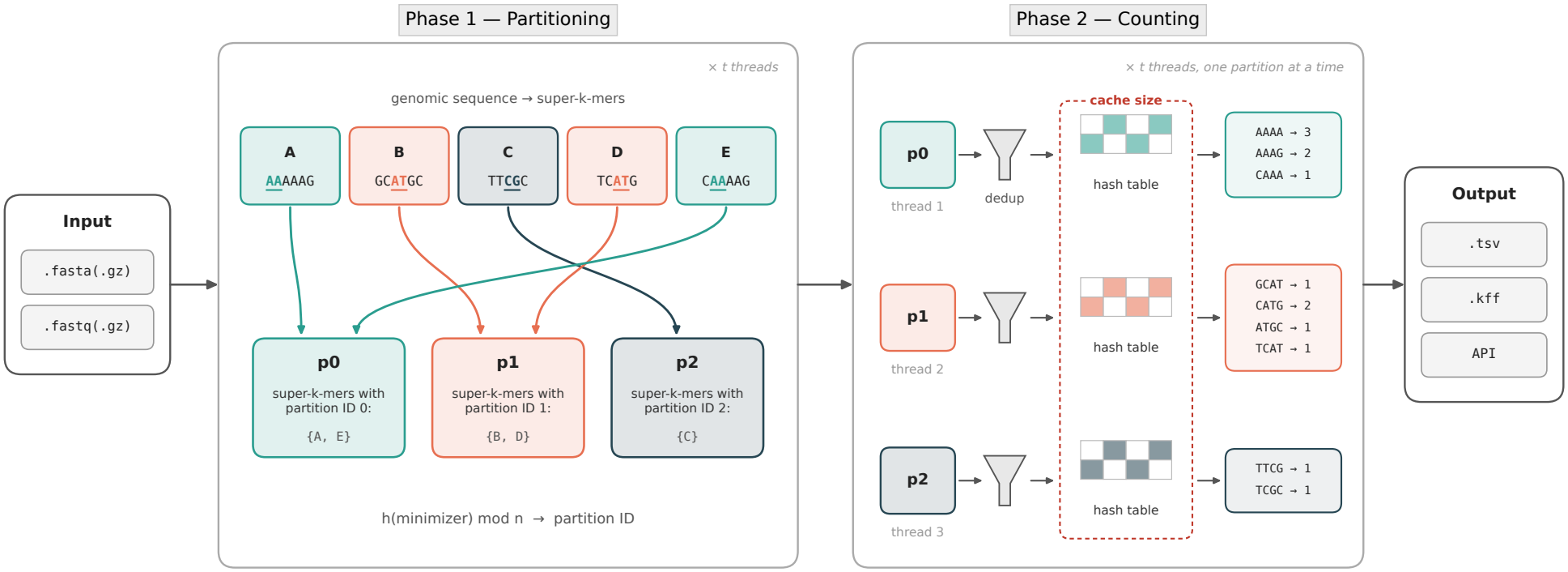
A schematic representation of the Tuna algorithm, for *k* = 4, *m* = 2, and with 3 threads, *n* = 3 partitions. For ease of illustration, non-canonical *k*-mers are counted. Phase 1 streams through the input files, computes super-*k*-mers from their genomic sequences, and routes each super-*k*-mer to a partition depending on its minimizer hash. For readability, only one stream is drawn, but Phase 1 is multi-threaded too, with several threads writing into the same partitions. During Phase 2, each thread processes one partition at a time, previously stored in memory or on disk, deduplicates the super-*k*-mers and inserts each super-*k*-mer’s *k*-mers into a hash table. *k*-mers are then emitted with their counts to the output. “API” in the “Output” box denotes programmatic access via a C++ callback, not a separate file format.

### 5.1. Phase 1: partitioning the input

#### Parsing

Phase 1 reads FASTA or FASTQ, plain or gzip-compressed, and turns it into a stream of bases packed two bits apiece. This is delegated to a C++ port of the Helicase library [Martayan et al., 2026], a vectorized parser for those formats. Ambiguous bases such as ‘N’ act as implicit sequence boundaries.

#### Thread scheduling

Thread scheduling in Phase 1 depends on whether the input consists of many files or a single large file. In the former case, each thread processes one file at a time, entirely, and then proceeds to the next file not yet assigned to any thread. A single compressed file does not allow this division of the work, since gzip decompression is inherently sequential. In this case, Tuna divides threads into two groups: the first group decompresses and parses the input, storing the resulting sequences in a queue of bounded size, while the second group retrieves the sequences from the queue and extracts their super-*k*-mers. When the queue is full, a thread from the first group also performs extraction until space becomes available, and once the whole input has been parsed, all threads perform extraction.

#### Routing by minimizer

Minimizers are computed with the two-stack sliding window minimum [Groot Koerkamp and Martayan, 2025], which takes amortized constant time per base. The order O is that of the random minimizer scheme of Section 2, applied to canonical *m*-mers: each *m*-mer and its reverse complement are encoded with two bits per base, and the smaller of the two values is hashed.

Consecutive super-*k*-mers are delimited by positions where the minimizer changes (Definition 2). Each super-*k*-mer is assigned to partition *p* = *h*(*µ*(*x*)) mod *n*, where *x* is any *k*-mer of the super-*k*-mer, *h* is a hash function, and *n* is the number of partitions. Hashing *µ*(*x*) is necessary because the hash value of a minimizer is the minimum over a window, so its high-order bits are biased toward zero: using this value directly would distribute the super-*k*-mers unevenly among the partitions, whereas applying *h* first yields a nearly uniform distribution.

Each super-*k*-mer is stored in its partition as a compact record, consisting of its length followed by its bases, encoded with two bits per base. Records are first accumulated in a buffer private to each thread and partition, and a full buffer is written to its partition at once, so that the lock of the partition is acquired only once per buffer.

#### Number of partitions

The number of partitions *n* sets how much input each one receives, and therefore how large the table built for it in Phase 2 will be. With *t* threads each holding one table during Phase 2, *n* decides what those *t* tables occupy together, which is precisely the miss(·) term of Section 4.3: at high abundance the analysis puts hashing up to one order of magnitude ahead of sorting when the tables are resident, and up to a factor of four ahead when they are not.

Tuna therefore chooses the smallest *n* for which the *t* tables in use at the same time are expected to fit in the last-level cache, whose size *M* is read at run time: 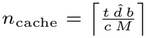, where 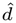 is an estimate of the number of distinct *k*-mers, *b* is the number of bytes a *k*-mer occupies in the table, and *c* = 3*/*4 leaves room for the operating system and the buffers of Phase 1.

Since 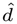 must be known before counting, it is estimated from the size of the input as 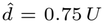, where *U* is the number of bytes of the uncompressed input, itself estimated as six times the size of a compressed file. The factor 0.75 is the fraction of bytes that are bases, the others being headers, line breaks, and ambiguous symbols; the estimate also assumes that the number of distinct *k*-mers equals the number of bases, which holds for genomic data at the lengths *k* of interest. This assumption fails for collections of closely related genomes, such as the assemblies of one species, which share most of their *k*-mers. For this reason, when the files of a collection are smaller than 50 MB on average, 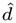 is computed from the largest file alone.

The final *n* is the larger of *n*_cache_ and one partition per 2 MB of input, so that the work can be balanced among the threads; it is then rounded up to a power of two, so that the modulo above reduces to a bit mask, and bounded by system limits. Section 6.1 shows that increasing *n* beyond this value can be counterproductive. Our heuristic for *n* rests on measurements made on our own datasets and may therefore carry their biases. Improving it and making it more robust is a natural direction for future work.

#### In-memory and on-disk pipelines

Most counters write the partitions to disk (Section 3), so that only the partition being counted resides in memory. The price is that the partitioned data is written and read once more in full. The analysis in Section 4 ignores this cost, since its two memory levels are the cache and the main memory, and rightly so: algorithms S and H would both pay it in the same measure. Tuna avoids it altogether whenever all the partitions fit in the memory budget, and writes them to disk only otherwise.

Tuna needs to know, before reading anything, roughly how much space the packed super-*k*-mers will take. It estimates this from the size of the input files in two steps: file bytes to bases, using a factor fixed by the format, since a FASTQ file spends about half its bytes on quality strings and compresses differently from a FASTA one, and then bases to super-*k*-mer bytes. Under random minimizers, the expected length of a super-*k*-mer is (*k* − *m* + 2)*/*2 *k*-mers, hence (3*k* − *m*)*/*2 bases, stored packed with a one-byte length header. Consecutive super-*k*-mers overlap by *k* − 1 bases, so each super-*k*-mer covers only (*k* − *m* + 2)*/*2 bases of the input, and the cost is 2(⌈(3*k* − *m*)*/*8⌉ + 1)*/*(*k* − *m* + 2) bytes per input base. At *k* = 31 and *m* = 17 that is 11 bytes per 8 bases, about 1.4 per base.

If the result sits below the budget, every partition is kept in memory and nothing touches intermediate storage. If it does not, Phase 1 streams each partition to its own file and Phase 2 memory-maps one partition per thread, so that peak memory is bounded by the write buffers while partitioning and by the live tables while counting. The estimate only decides which pipeline to start in. Should it prove optimistic, the in-memory buffers spill to disk on crossing the budget and the run continues exactly as if it had started there.

### 5.2. Phase 2: counting a partition

#### Deduplicating super-*k*-mers

Deduplication is the source of the advantage of H in Section 4: it makes the counting work proportional to *N*_*d*_, the number of *k*-mer occurrences in the distinct super-*k*-mers of a partition, rather than to *N*, the total number of *k*-mer occurrences in the partition. Without deduplication, a super-*k*-mer occurring *r* times would require *r* insertions of each of its *k*-mers. Tuna therefore first replaces the identical super-*k*-mers of a partition with a single entry storing their multiplicity, then inserts the *k*-mers of each distinct super-*k*-mer once, incrementing their counters by that multiplicity.

The difference between *N* and *N*_*d*_ reflects the redundancy of the input: the saving is large on high-coverage reads and negligible on assemblies, where the additional pass costs more than it saves. Tuna therefore examines a sample of the first records of each partition, and skips deduplication when repeated super-*k*-mers are too rare, or when the auxiliary table would exceed its share of the memory budget.

#### Hash tables

Each partition is counted in its own table, so the *t* threads never synchronize while counting. The *k*-mers of a super-*k*-mer are computed incrementally, each from the previous one by shifting in one base, and are inserted, in canonical form, into a hash table with open addressing and robin-hood probing [Leitner, 2022].

#### Sizing a table

The number of slots of a table should be proportional to the number of distinct *k*-mers of the partition, which is unknown until the partition has been counted. The only quantity available beforehand is the number of *k*-mer occurrences, known from Phase 1. This is an upper bound, attained only if no *k*-mer repeats, and sizing every table from it would waste the cache space that the choice of *n* is meant to preserve. Tuna therefore uses this bound only for the first partitions: once a partition has been counted, its number of distinct *k*-mers is used to size all the partitions counted afterwards.

#### Output

Once all *k*-mers from a partition are inserted in their table, Tuna scans it and outputs the pairs (*k*-mer, count) whose counts lie within the thresholds of Problem 1. The output is either a plain TSV file, or a file in the binary KFF format [Dufresne et al., 2022]. Tuna also provides a C++ library interface, through which another tool can run the pipeline and receive the pairs through a callback function, without writing any file.

## 6. Results

In this section, we validate the design of Tuna experimentally and compare it against the state of the art, KMC3 (v3.2.4) [Kokot et al., 2017c] and FastK (v1.2) [Myers, 2021]. FastK is included despite the absence of a peer-reviewed publication because, as already mentioned, it is in routine use in assembly pipelines (see references in Section 3). The hash-based counters reviewed in Section 3 are not included instead: they were designed before the disk-partitioned organization became standard and are reported as slower than KMC3 in the published comparisons [Kokot et al., 2017c; Erbert et al., 2017], so KMC3 and FastK are the appropriate reference points.

Tuna is written in C++ and is available, together with reproducibility scripts, at github.com/vicLeva/tuna.

### Methodology

For all experiments reported here, we use version 1.5.0 of Tuna. All tested software was compiled with gcc 14.2.0 and flags `-O3 -march=native`. We report the total wall-clock time (including the I/O time to read the input and write the output), peak resident memory, and the peak volume of temporary data a tool holds on disk. All experiments count canonical *k*-mers. Unless stated otherwise, every experiment uses *k* = 31 (we also use *k* = 27 in some experiments).

All tools are tested with their default parameters. The only settings we fix are common to the three: the *k*-mer length, the thread count (8), the memory budget (256 GB), and the reporting of every *k*-mer that appears at least once (i.e., we set *c*_min_ = 1 and *c*_max_ high enough that no tool saturates any count). We verified that the three tools report identical sets of (*k*-mer, count) pairs on every input. Tuna also requires a minimizer length: we set *m* = 13 for *k <* 31 and *m* = 17 for *k* ≥ 31, except for *k <* 15 where *m* = *k* − 2. In the experiments that follow, we therefore use *m* = 13 for *k* = 27 and *m* = 17 for *k* = 31.

Every run writes the tool’s own compact binary format (KFF [Dufresne et al., 2022] for Tuna).

### Hardware

All experiments ran on a GenOuest node with two 32-core Intel Xeon Gold 6338N processors at 2.20 GHz, 64 cores in all, each socket holding 48 MB of last-level cache, and 1 TB of memory, under Debian GNU/Linux 12. Storage is an array of SATA SSDs in RAID 6, measured at 520 MiB/s sequential read and 480 MiB/s sequential write.

### Datasets

We use the eight collections summarized in Table 1, spanning assemblies and sequencing reads, single species and metagenomes. They range from single samples to pangenome-scale cohorts of twenty-five thousand genomes. They cover bacterial, chicken, and human genomes, alongside gut and sea water metagenomes. Their size runs from under a megabyte per sample to more than six hundred gigabytes for a whole collection. Several of them were used in the KMC3 evaluations, which keeps our measurements comparable with the published ones. Links to data access are provided on the Tuna GitHub repository.

**Table 1.** Dataset statistics. The last three columns are medians over the samples, each counted on its own. The abundance of a sample is its number of *k*-mer occurrences divided by its number of distinct *k*-mers (written *c* in Section 4): it stays close to 1 on assemblies and grows with sequencing depth. All inputs are gzip-compressed files: FASTA for assemblies and FASTQ for reads.

| Dataset | Organism | Type | Files | Size<br>per sample | Distinct *k*-mers<br>per sample | Abundance<br>per sample |
|---|---|---|---|---|---|---|
| Ecoli | *E. coli* | assemblies | 3500 | 1.6 MB | 5.04 M | 1.02 |
| Salmonella | *S. enterica* | assemblies | 25000 | 1.5 MB | 4.90 M | 1.00 |
| Gut | gut MAGs | assemblies | 10000 | 0.6 MB | 2.10 M | 1.01 |
| Human | *H. sapiens* | assemblies | 60 | 0.9 GB | 2.50 G | 1.22 |
| Tara | Tara Oceans (sea water) | reads | 12 | 2.9 GB | 2.05 G | 1.27 |
| Gallus | *G. gallus* | reads | 12 | 2.1 GB | 0.97 G | 2.21 |
| Human3 | *H. sapiens* | reads | 36 | 17.1 GB | 3.02 G | 4.65 |
| HumanR | *H. sapiens* | reads | 1 | 2.5 GB | 2.86 G | 2.25 |

### 6.1 Validation

The experiments in this section serve two purposes: to validate the number of partitions automatically chosen by Tuna, which we then keep in all subsequent experiments, and to test whether the analysis of Section 4 correctly predicts that sorting and hashing separate as the abundance of the *k*-mers grows.

#### Effect of the partition count

The number of partitions *n* is chosen by Tuna at run time (Section 5.1). Figure 2 measures its effect: Gallus and Human3, the whole collections of 12 and 36 read files, are counted repeatedly with nothing changed but *n*, doubled from 2^5^ to 2^21^ on Gallus, and to 2^18^ on Human3^2^.

**Fig. 2.**
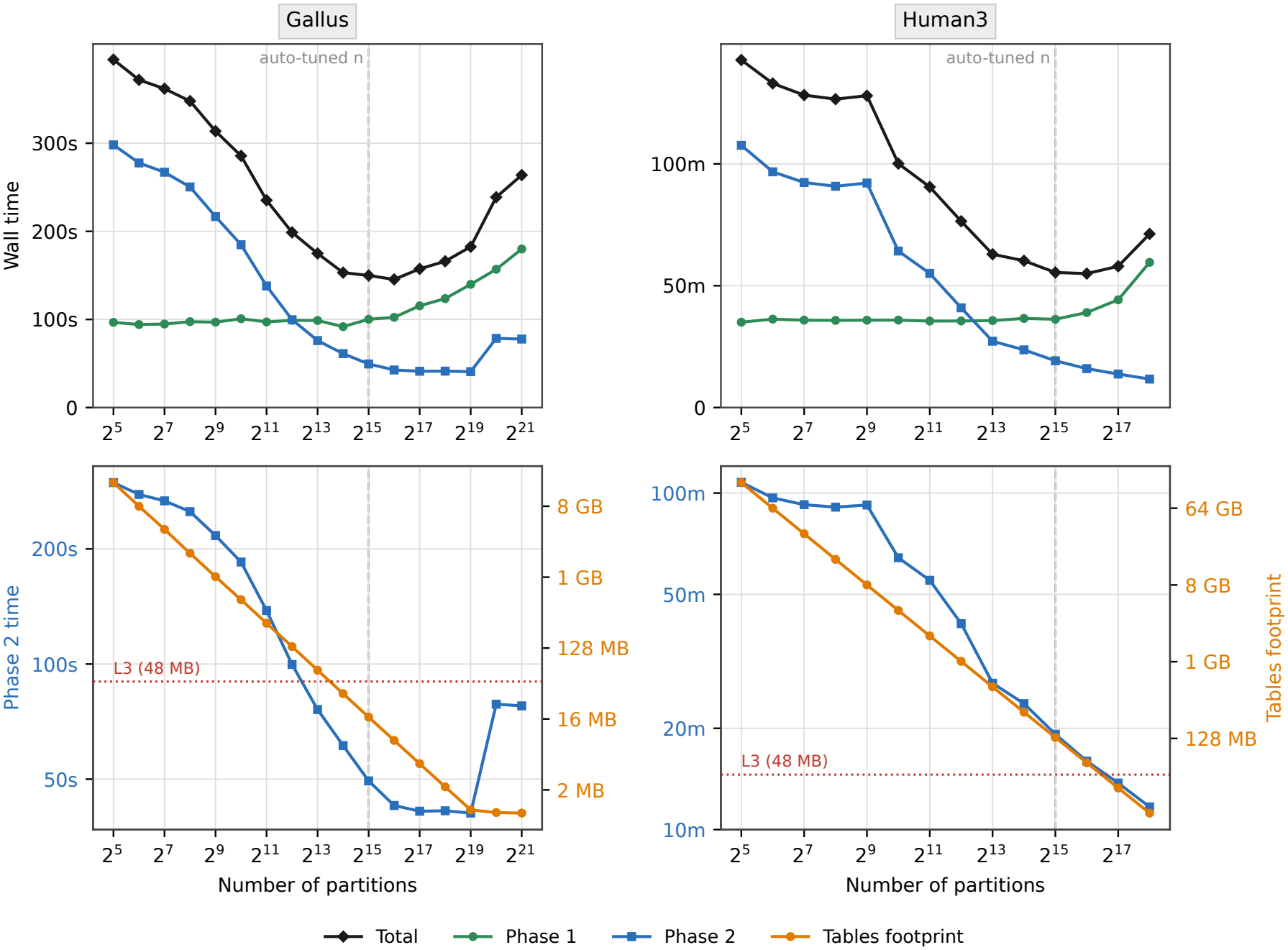
Gallus and Human3 counted repeatedly with only the number of partitions *n* changed (*k* = 31, *m* = 17, 8 threads). Top: wall-clock time and its two phases. Bottom: Phase 2 time against the combined footprint of the 8 tables held at once; the dotted horizontal line marks the 48 MB L3 cache. In all panels, the dashed vertical line marks the value of *n* that Tuna chooses automatically.

The two phases pull in opposite directions. Partitioning (Phase 1) is indifferent to *n* while partitions remain few, and only begins to cost once managing that many of them at the same time becomes expensive in itself, beyond about 60,000 here. Counting (Phase 2) moves the other way: a larger *n* spreads the distinct *k*-mers over more partitions, so the *t* tables held at once shrink and their insertions miss the cache less often. The lower panels show this, plotting Phase 2 time against the combined footprint of those tables. On Gallus Phase 2 falls steeply while that footprint exceeds the last-level (L3) cache shared by the threads, and keeps falling past the crossing until each of the *t* tables is about a megabyte, which happens to be the size of one core’s L2, after which it flattens. The best total time sits there, at *n* = 2^16^, where Phase 1 in turn begins to rise. Human3 holds seven times more distinct *k*-mers, so bringing its tables that low would cost more in partitioning than it returns, and its optimum stays above the L3 line, at a footprint of 72 MB. Past the optimum a finer split rewards neither phase, and on Gallus even Phase 2 itself eventually turns back up. The same upturn is expected on Human3, at a partition count above the range the sweep could reach there.

The auto-tuning of Section 5.1 lands one power of two below the optimum of the sweep on both collections, within 3.7% of the best time observed on Gallus and within 2.1% on Human3. We therefore keep that choice, which is Tuna’s default, in every other experiment.

#### High abundance

Section 4 predicts that the two families separate as *k*-mer abundance grows, the cost of sorting following the number of *k*-mer occurrences and that of hashing the number of distinct ones. Figure 3 tests this directly. HumanR, a single read file of 2.5 GB, is counted at increasing coverage, from 1× to 100×. Up to 5× each level is a subsample of the original file, so the distinct set grows with it. Beyond that the reads are replicated, which freezes the distinct set near 2.9 billion while occurrences rise by a factor of twenty, isolating abundance from diversity.

**Fig. 3.**
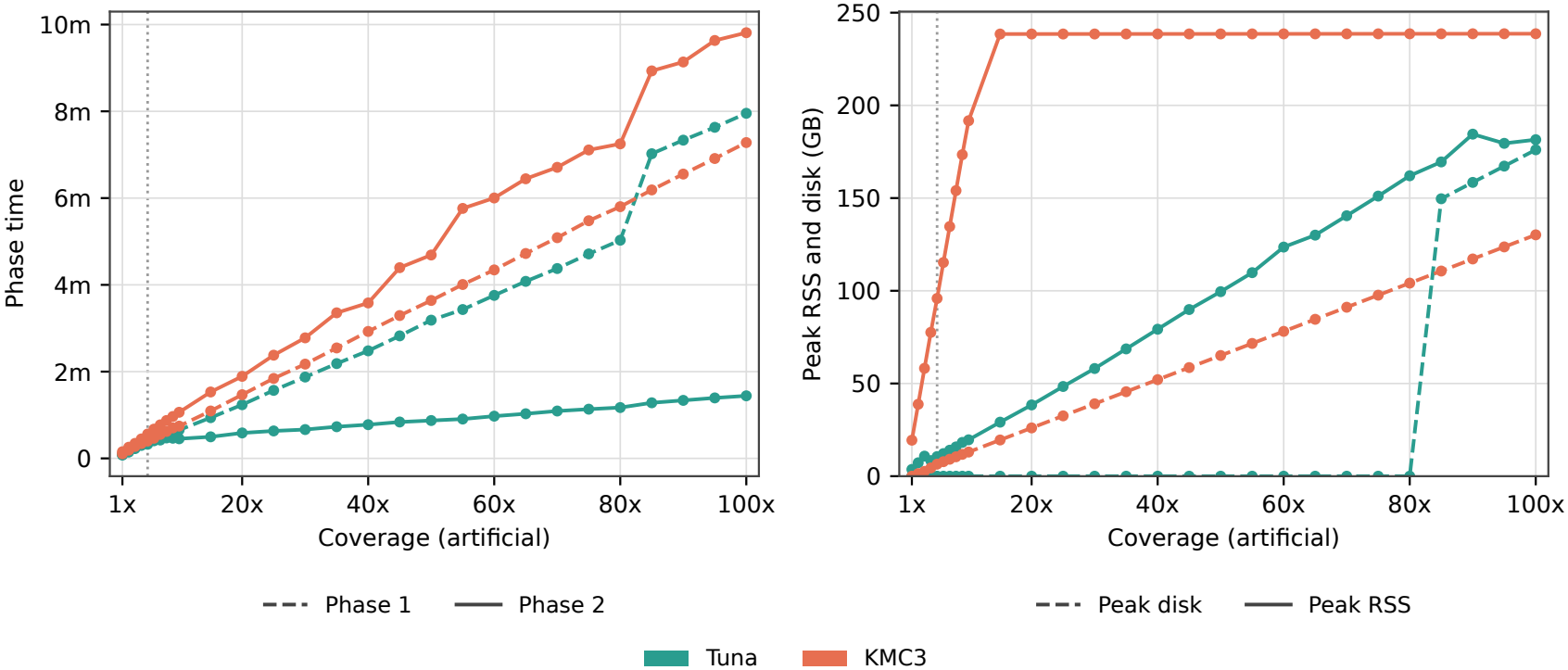
HumanR counted at increasing coverage, from 1*×* to 100*×* (*k* = 31, *m* = 17, 8 threads). Left: the two phases timed separately. Phase 1 (dashed) is the same algorithm in both tools, so the curves nearly coincide; Phase 2 (solid) is where sorting and hashing differ, and the curves separate. Right: peak resident memory (solid) and peak temporary data on disk (dashed). The dotted vertical line marks 5*×*, the coverage of the source read set: below it each level is a subsample and the distinct *k*-mer set grows with coverage; above it the reads are replicated and only the abundance grows.

The measurements closely follow the analysis. First of all, partitioning costs the two tools much the same, Tuna staying between 1.1 and 1.2× ahead up to 80×, so the separation comes entirely from counting (Phase 2). There, the counting time of Tuna only rises by a factor of 19 across the sweep whereas that of KMC3 rises by a factor of 64, widening the gap on that phase alone from 2.0 to 6.8×. (Since partitioning takes the larger share of Tuna’s time, the gain over a whole run stays smaller than on counting alone.) It grows from 1.6 to a peak of 2.2× at 55×, then falls back to 1.8× once the partitioned data outgrows the memory budget and Tuna moves to its disk pipeline (Section 5.1), which costs it about 40% more partitioning time and accounts for the temporary data in the right panel. The analysis predicts a factor between four and ten at high abundance, depending on whether the tables stay in cache. The 6.8× measured on Phase 2 falls inside that range.

### 6.2. Performance

The experiments in this section model the two scenarios in which *k*-mer counting is used in practice. In the first, each sample is counted on its own, as in genome assembly, assembly quality assessment, and genome profiling [Rhie et al., 2020]: what matters is the time per file, which compounds over collections of thousands. In the second, a single count database is built over a whole collection, as when constructing indexes for sequence search: what matters is the cost of one large run. We cover the single-file scenario first, then the whole-collection one, counting each of the five smaller collections in a single run under the default configuration, and finally stress-test the latter scenario on the two largest read collections, Gallus and Human3, where we also vary the *k*-mer length and the memory budget.

#### Single file

Figure 4 shows the median computation time per file for the three tested tools. Counted one file at a time, Tuna has the lowest median wall time on every dataset: this is aligned with the analysis in Section 4 because the hash tables always fit in cache for these experiments. In particular, against KMC3 the speedup factor runs from 1.1 to 3.7×, and against FastK from 1.7 to 3.8×. Memory follows the same trend once the inputs are large. On the three bacterial collections, all three tools sit near a hundred megabytes and differ by a few tens either way, which is noise at that scale. On Human and Tara, where a single file holds billions of distinct *k*-mers, Tuna uses 5.2 and 4.2 GB where KMC3 uses 45.2 and 37 GB, and FastK 32 and 29.4 GB.

**Fig. 4.**
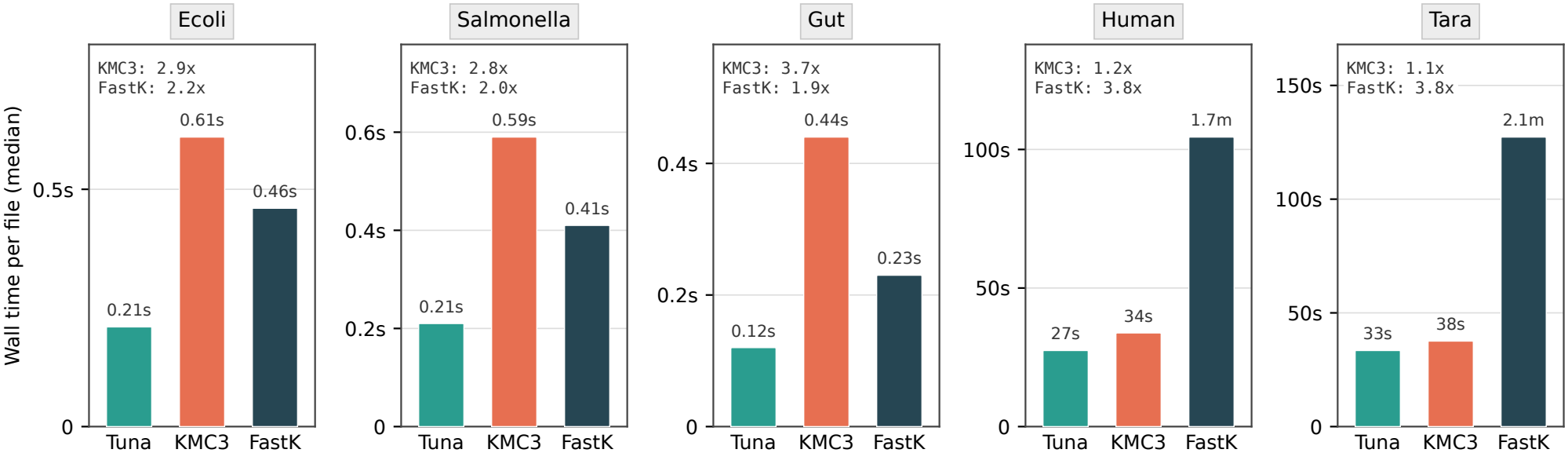
Median wall-clock time per file when each sample is counted on its own (*k* = 31, *m* = 17, 8 threads). The annotations over the bars give the speedup of Tuna over each competitor.

#### Whole collection

Figure 5 shows the wall-clock time when counting whole collections. Tuna is the fastest of the three on four of the five, ahead of KMC3 by 4.3× on Ecoli, 3.4× on Human, 3.0× on Salmonella and 1.6× on Gut, and ahead of FastK by 1.6 to 2.7× wherever it completes. FastK fails on Salmonella and exceeds the time limit on Tara.

**Fig. 5.**
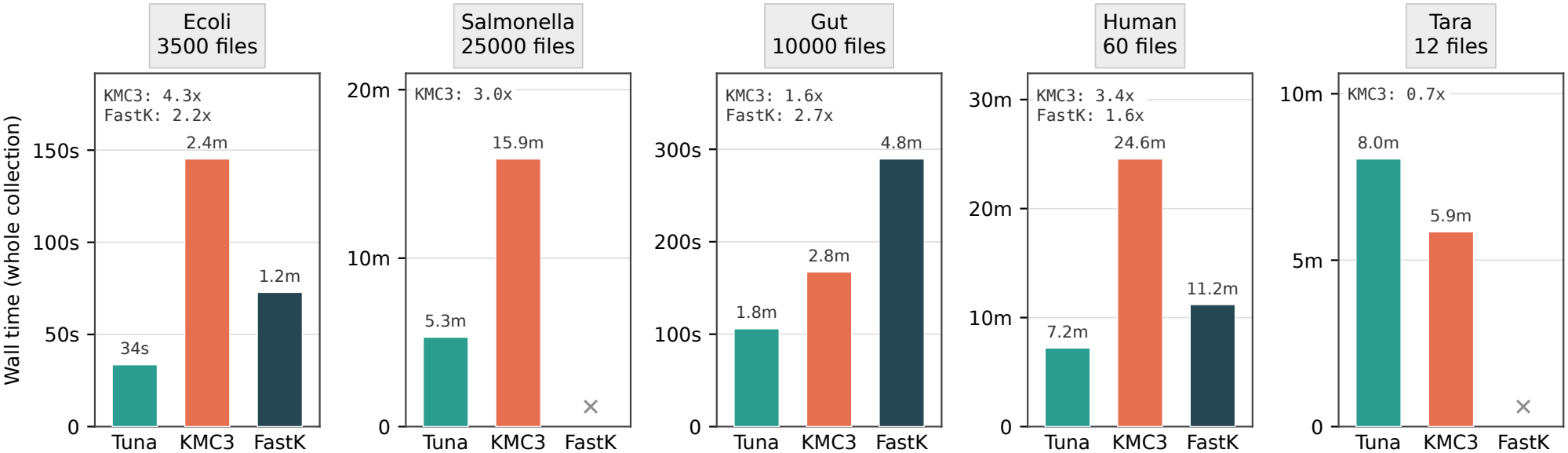
Wall-clock time for each collection counted in a single run over all of its files (*k* = 31, *m* = 17, 8 threads). The annotations over the bars give the speedup of Tuna over each competitor. A cross marks a run that produced no result: FastK failed on Salmonella and exceeded the 6-hour time limit on Tara.

Tara is the exception, Tuna taking 8.0 minutes against 5.9 for KMC3 (36% more). It is also the only collection that meets both conditions Section 4 gives for sorting to prevail (low abundance and hash tables out of cache). Its 12 read sets hold 22.5 billion distinct *k*-mers at an abundance of 1.6, so the data is almost free of repetitions. At the 2^15^ partitions Tuna selects, each table holds some 690 thousand distinct *k*-mers, and the eight tables live at once occupy about 140 MB against the 48 MB of the L3 cache. The same estimate puts the other four between 2 and 41 MB, inside the cache or at its edge. Phase 2 follows that ordering, Tuna leading by 13.2 and 14.5× on Salmonella and Ecoli at abundances of 216 and 109, by 7.2× on Human at 59, by 1.5× on Gut at 3.0 with its tables at 41 MB, and trailing by 1.6× on Tara. End-to-end the factors are smaller throughout, Phase 1 taking most of the run at this scale.

KMC3 fills the budget it is given on every collection, about 238 GB, where Tuna peaks between 4.8 and 62 GB. What Tuna writes to disk follows the pipeline it picks: nothing at all on Ecoli, Gut and Tara, which it counts in memory, against 154 and 230 GB on Salmonella and Human, where it switches to the disk pipeline. KMC3 writes between 16 and 164 GB.

#### Stress test on large read collections

We count here the whole Gallus and Human3 collections in a single run, 12 and 36 read files amounting to 24.0 and 614 GB of compressed input, and compare Tuna against its competitors in Table 2. The 8 GB budget is the memory a laptop or a modest workstation offers, and *k* = 27 is the second length the analysis instantiates alongside *k* = 31. These two values per parameter are enough to expose the behaviors of interest, a second *k*-mer length activating the (*k, x*)-mer machinery of the sort and a tight budget forcing the disk pipelines. Both collections are read sets. Counted as single collections they average 8.5 and 24.8 occurrences per distinct *k*-mer, well above the per-sample figures of Table 1, and it is that regime where the analysis expects hashing to win.

**Table 2.** Gallus and Human3, each counted in a single run over its whole set of files, at two *k*-mer lengths (*m* = 13 at *k* = 27, *m* = 17 at *k* = 31) and under two memory budgets (8 threads). The budget is the value passed to each tool. Bold marks the best time in each column. FastK did not complete within the time limit on Gallus, and it reports no per-phase timings, so its Phase 1 and Phase 2 entries are marked “n/a.”

| Dataset | RAM budget (GB) | Tool | *k* = 27 |  |  |  |  | *k* = 31 |  |  |  |  |
|---|---|---|---|---|---|---|---|---|---|---|---|---|
|  |  |  | Time (min) |  |  | Peak RSS (GB) | Peak disk (GB) | Time (min) |  |  | Peak RSS (GB) | Peak disk (GB) |
|  |  |  | Phase 1 | Phase 2 | Total |  |  | Phase 1 | Phase 2 | Total |  |  |
| Gallus | 8 | Tuna | **1.9** | **0.8** | **2.8** | 2.1 | 32.0 | 2.0 | **0.9** | **2.9** | 2.1 | 33.3 |
|  |  | KMC3 | 2.1 | 1.8 | 3.9 | 7.5 | 26.0 | **1.5** | 2.1 | 3.6 | 7.5 | 23.9 |
|  | 256 | Tuna | **1.6** | **0.8** | **2.4** | 33.6 | 0.0 | **1.6** | **0.8** | **2.5** | 34.9 | 0.0 |
|  |  | KMC3 | 2.4 | 1.5 | 3.9 | 183.6 | 26.8 | 1.7 | 1.8 | 3.5 | 238.5 | 24.2 |
| Human3 | 8 | Tuna | **36.6** | **18.5** | **55.2** | 3.1 | 679.4 | 37.5 | **18.7** | **56.2** | 2.6 | 707.2 |
|  |  | KMC3 | 43.6 | 32.1 | 75.7 | 23.0 | 571.5 | **29.6** | 70.0 | 99.6 | 29.5 | 522.9 |
|  |  | FastK | n/a | n/a | 91.5 | 7.6 | 932.2 | n/a | n/a | 81.8 | 7.6 | 768.8 |
|  | 256 | Tuna | **36.4** | **18.6** | **54.9** | 3.7 | 679.4 | 40.4 | **18.6** | **59.0** | 3.8 | 707.2 |
|  |  | KMC3 | 46.1 | 28.0 | 74.1 | 239.0 | 572.8 | **32.8** | 34.3 | 67.1 | 238.9 | 522.9 |
|  |  | FastK | n/a | n/a | 70.5 | 168.2 | 540.3 | n/a | n/a | 71.2 | 192.1 | 509.9 |

Tuna has the shortest total time of the three tools in every configuration of Table 2. FastK completes only Human3, none of its Gallus runs finishing within the six-hour limit, and there it trails Tuna in all four configurations, by between 1.2 and 1.7×. FastK reports no phase split, so the phase-by-phase discussion below concerns Tuna and KMC3 only.

The analysis covers Phase 2, the one phase in which the two strategies differ. Partitioning takes about two thirds of the run at this scale, so the end-to-end margins are correspondingly smaller than the Phase 2 figures. Phase 1 is the same algorithm in the two, and at *k* = 31 KMC3 is the faster of the two in three of the four runs, by up to 1.3×, whereas at *k* = 27 Tuna is ahead in all four, by between 1.1 and 1.5×. The separation therefore comes entirely from Phase 2, where Tuna leads in every configuration, at *k* = 31 by between 2.2 and 2.4× on Gallus and between 1.9 and 3.7× on Human3. That separation narrows at *k* = 27, to between 2.0 and 2.2× on Gallus and to between 1.5 and 1.7× on Human3, which is what Section 4 predicts. At *k* = 27 the sort operates on (*k, x*)-mers with *x* = 3, so a record carries up to four *k*-mer occurrences and the merge term decays as the abundance grows, bringing the modelled cost of sorting to about 4.5 ns per occurrence at an abundance of 20, against 13.5 at *k* = 31, where *x* is zero and no merge takes place. The cost of hashing does not depend on *k*. Sorting thus recovers part of the gap at *k* = 27 and gives it back at *k* = 31, clearly on Human3 and only marginally on Gallus.

Setting the two memory budgets side by side shows how each tool behaves when memory is scarce. Tuna counts Human3 through its disk pipeline under either budget, so its time barely moves at *k* = 31, 56.2 minutes under the 8 GB budget against 59.0 under the full one. On Gallus the tighter budget carries it from the in-memory pipeline to the disk one (Section 5.1), peak memory falling from 34.9 to 2.1 GB for 0.4 minutes of extra time. KMC3 instead fills whatever memory it is given, and pays when there is less of it: on Human3 at *k* = 31 it takes 67.1 minutes under the full budget and 99.6 under 8 GB, nearly all of the difference falling in Phase 2, which goes from 34.3 to 70.0. FastK moves the same way more gently, 71.2 minutes under the full budget against 81.8 under 8 GB. Gallus is small enough that the restriction costs either competitor nothing. We note that KMC3 does not hold strictly to the budget it is given, peaking at 29.5 GB when asked for 8. The budget applies to memory alone. The last column of Table 2 reports the temporary data each tool writes to disk. On Human3 all three write hundreds of gigabytes under either budget, 707 GB for Tuna, 523 for KMC3 and between 510 and 769 for FastK. On Gallus with the full budget Tuna writes nothing at all, the run staying in memory from end to end, where KMC3 still writes 24 GB, and under the 8 GB budget Tuna writes 33 GB.

## 7. Conclusion

We set out to settle whether sorting or hashing is the better way to count the *k*-mers in a set of super-*k*-mers. Our analysis indicates that the answer depends on two factors: how abundant the *k*-mers are and whether the tables stay in cache. In particular, hashing is faster than sorting whenever the hash tables fit in cache. Even when the tables do not fit in cache, hashing beats sorting once the *k*-mers are abundant. The only case in which sorting may be faster than hashing, according to the analysis, is when the hash tables do not fit in cache and the *k*-mers are mostly unique. We observed this once, on the Tara dataset, the single input in our evaluation where Tuna is not the fastest algorithm, for the reasons discussed in Section 6.2. No other collection we tried meets both conditions, and they are in tension. Tables large enough to leave the cache require a large distinct set, and at that scale the data providing one is sequencing reads, which are usually redundant. Assemblies stay close to unique but remain small enough for the partitions to keep the tables resident. A seawater metagenome, like Tara, is one of the few places the two meet, its diversity being large enough that sequencing more returns new *k*-mers rather than repeats of those already seen. The reversal in performance for this case is not an anomaly but the analysis holding where it predicts the other outcome.

To validate the analysis, we implemented a hashing-based algorithm called Tuna. We show that Tuna outperforms KMC3 on all but one input by up to 4.3×, with the margin widening to 6.8× on the counting phase as abundance grows, in accordance with the analysis.

Two directions seem promising for future work. First, a hybrid algorithm could choose between hashing and sorting for each partition, based on its estimated abundance and table size, removing the only case in which Tuna is slower than KMC3. Second, deduplicating *canonical* super-*k*-mers could nearly double the effectiveness of deduplication on sequencing reads, where about half of the occurrences of a super-*k*-mer come from the opposite strand.

## Funding

This work was supported by Inria challenge program OmicFinder.

## Competing interests

R.P. is a co-founder of Ocean Genomics Inc.

## Footnotes

1 We omit ceilings and floors when they are not necessary.

2 The sweep stops earlier on Human3 because its disk pipeline keeps one file open per partition and so reaches the system limit on open file descriptors.

## Notes

https://github.com/vicLeva/tuna

